# Adapting CRISPR-associated transposons for rapid and high-throughput reverse genetics

**DOI:** 10.1101/2025.10.31.685825

**Authors:** David W. Basta, Franz G. Zingl, Yiyan Yang, Kaylee T. Nguyen, Yang-Yu Liu, Matthew K. Waldor

## Abstract

CRISPR-associated transposons (CAST) use guide RNAs to direct their transposition and are being harnessed as tools for programmable genome engineering across diverse bacterial species. However, CAST systems have not been adapted for high-throughput genetic screening. Here, we present MultiCAST, a streamlined platform for rapid and scalable guide RNA-directed transposon insertion in bacteria. MultiCAST generates targeted insertions in a single step through conjugative delivery of conditionally replicative plasmids encoding the CAST enzymatic machinery and a selectable mini-transposon expressing the guide RNA. By leveraging the inserted guide sequence as a molecular barcode, MultiCAST enables pooled, high-throughput genetic screens using only amplicon sequencing. We identified factors that influence transposition efficiency and the accuracy of insertion frequency measurements derived from guide sequencing. Adjusting the ratio of donor and recipient strain during conjugation mitigates guide-transposon “crosstalk”, in which a single recipient cell acquires multiple donor plasmids containing distinct guides. Furthermore, we developed a machine learning-based predictive model for selecting highly active guides based on target sequence features that strongly correlate with activity. The nucleoid-associated protein H-NS was also found to inhibit CAST activity, providing a mechanistic explanation for variable insertion frequencies among non-essential genes. To demonstrate the scalability of MultiCAST, we screened a pooled mutant population created from >5,200 guides targeting 88 genes in *E. coli* across twelve nutrient conditions, accurately identifying genes with condition-specific fitness effects. The simplicity, speed, and throughput of MultiCAST make genome-scale functional screens more accessible across a wide range of bacterial species.

**Significance:** Efficient gene disruption is essential for understanding bacterial gene function, but traditional genetic approaches are labor-intensive and generally not well-suited for high-throughput studies. We developed MultiCAST, a simple and scalable method that harnesses guide RNA-directed CRISPR-associated transposons for targeted bacterial gene disruption. MultiCAST enables single and pooled transposon mutagenesis in a single step and eliminates the need for complex sequencing library preparation protocols by using the guide sequence as a quantifiable surrogate for mutant abundance. This approach allows thousands of mutants to be generated and screened simultaneously across multiple conditions using only amplicon sequencing. By dramatically reducing the time, cost, and complexity of reverse genetics, MultiCAST opens new possibilities for genome-scale functional studies, accelerating the discovery of bacterial gene functions.

## Introduction

Transposon mutagenesis is a foundational method in bacterial genetics (1). Identification of the genomic location of randomly inserted transposons facilitates understanding of causal links between genotypes and phenotypes. The pairing of next-generation sequencing (NGS) with transposon mutagenesis (Tn-seq) has made it possible to identify the insertion location and relative abundance of thousands of transposon mutants in a pooled population, enabling genome-scale assessment of the genetic determinants underlying a given phenotype. However, Tn-seq requires a multi-step protocol involving multiple days of sample preparation, numerous reagents, and complex computational pipelines in order to directly map the transposon-genome junction, creating ample opportunity for cascading errors during sample preparation and analysis (2).

We hypothesized that CRISPR-associated transposons (CAST) could serve as a simple alternative to the experimental and computational workflows of traditional Tn-seq screens. CAST use RNA-guided CRISPR-Cas systems to direct their transposition (3–5). The CRISPR-Cas complex identifies a target DNA sequence (protospacer) via a CRISPR RNA (crRNA or guide) and recruits Tn7-like transposition enzymes to insert the transposon downstream of the target site (Fig. 1a). The programmable nature of the guide RNA has made CAST systems potent tools for genome engineering in both bacterial and eukaryotic cells, where they enable targeted insertion of kilobase-scale DNA payloads (6–13), and for bacterial functional genetics, where they enable targeted gene inactivation (7, 14–18).

**Fig. 1.**
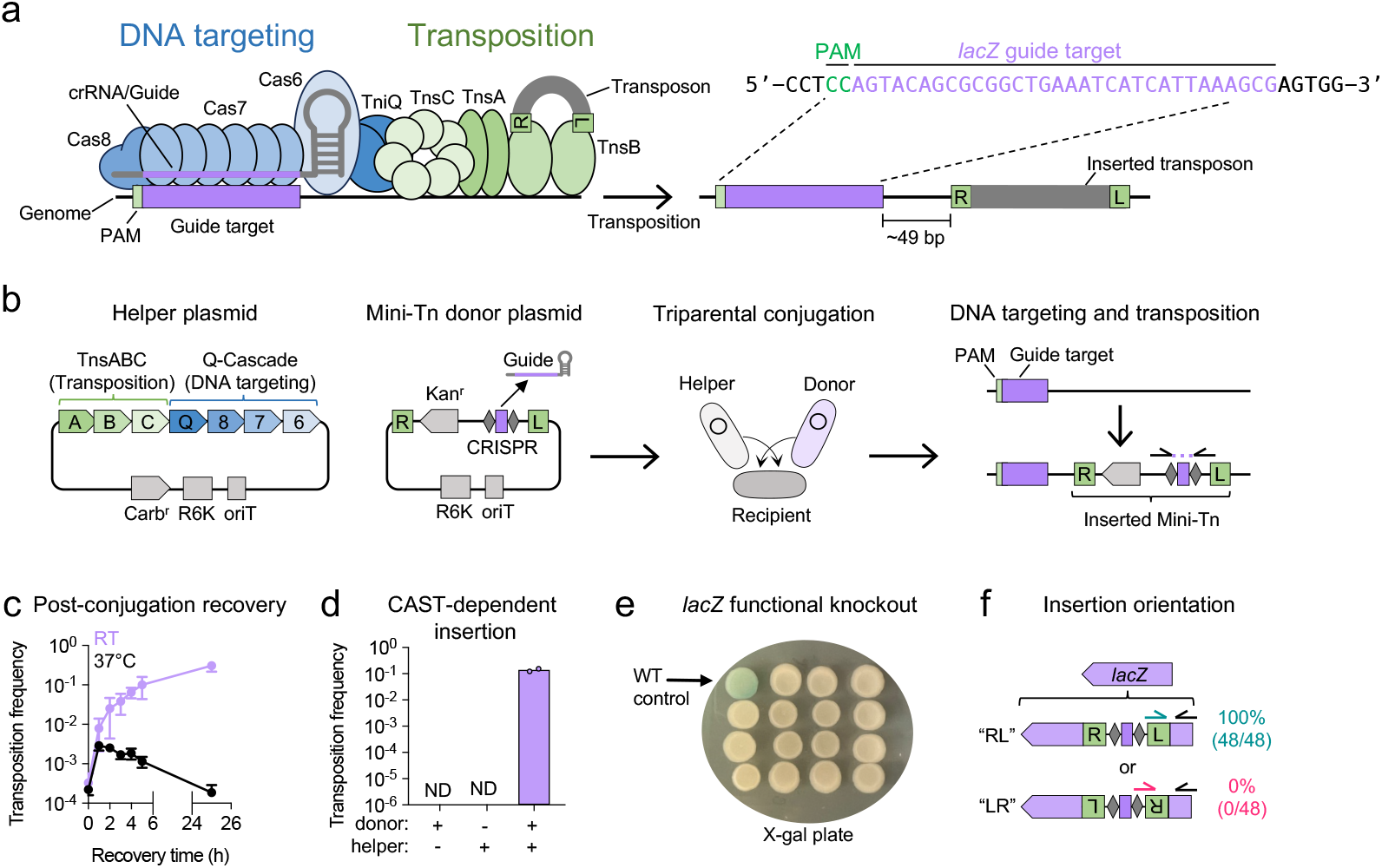
Single-gene knockout using MultiCAST. **a)** Illustration of components of Type I-F3 CAST system. The 32-nt CRISPR RNA (crRNA/guide) is flanked by repeat sequences (gray diamonds) recognized by the Cascade DNA targeting complex. The guide-Cascade complex binds a target site adjacent to a ‘CN’ protospacer adjacent motif (PAM) and directs insertion of the transposon ~49 bp downstream. ‘R’ and ‘L’ denote the right and left transposon ends. **b)** Overview of the MultiCAST platform and workflow. The CAST machinery is encoded on a helper plasmid, while the mini-Tn donor plasmid contains an antibiotic selectable marker flanked by transposon ends. A guide oligonucleotide is assembled into the linearized donor plasmid and transformed into a conjugative donor strain (MFD*pir*). Triparental conjugation is then performed with the donor, helper, and recipient strains. The CRISPR and transposase machinery are supplied by the helper plasmid, while the guide RNA expressed from the donor directs site-specific transposition. The inserted guide is amplified and sequenced to quantify mutant abundance. Kan^r^ = kanamycin resistance gene. Carb^r^ = carbenicillin resistance gene. oriT = RP4 origin of transfer. R6K = π-dependent origin of replication. **c)** Transposition frequency following post-conjugation recovery at room temperature (RT) vs 37 °C. Frequency is calculated as the ratio of antibiotic-resistant CFU to total CFU. Data represent mean ± S.D. of three biological replicates. **d)** Control experiment confirming that transposition requires both donor and helper plasmids. ND = Not detected. **e)** Functional validation of *lacZ* disruption. Randomly picked transconjugants were spotted onto LB agar containing X-gal. Loss of β-galactosidase activity results in white colonies. **f)** Orientation-specific PCR showing that transposon insertions predominantly occur in the ‘RL’ orientation (right end 5′ to left end). Arrows indicate PCR primer positions.

An appealing property of CAST systems is that the designed guide sequence itself can function as an intrinsic barcode, enabling quantification of mutant abundance via amplicon sequencing. This method uses a single PCR reaction to amplify the barcode sequence encoded within each transposon (19), and the NGS read count of each barcode then serves as a proxy for the abundance of each mutant. Compared to Tn-seq, amplicon sequencing is faster, more cost-effective, and scalable across multiple test conditions. Although CAST systems have recently been adapted for genome-scale functional screens (20, 21), the accuracy of guide-based amplicon sequencing as a proxy for CAST mutant abundance has not yet been benchmarked against Tn-seq, which represents the “ground truth” for transposon insertion frequency.

Moreover, fitness-independent factors that influence CAST insertion are not well described. For example, the frequency of each transposon insertion in a pooled mutant population is typically interpreted as a proxy for mutant fitness (22). However, this interpretation can be confounded by steric hindrance from DNA-binding proteins such as the nucleoid-associated protein H-NS, which may block transposition into non-essential genes and lead to their misclassification as essential (23–25). In CAST systems, variability in guide activity can also distort fitness estimates (21). Therefore, characterization of non-fitness-linked factors influencing CAST insertion is critical for the design and interpretation of large-scale CAST mutagenesis screens.

To address these challenges, we adapted the Type I-F3 CAST Tn6677 from *Vibrio cholerae* strain HE-45 (4, 7) into a streamlined platform called MultiCAST for bacterial genome engineering and functional genetics. Type I-F CAST systems have been adapted for use in both bacteria and eukaryotes due to their high targeting specificity (7, 12, 13, 26). In MultiCAST, the Tn6677 CRISPR-Cas and transposition proteins are encoded on a helper plasmid, while the guide RNA is cloned into a selectable mini-transposon (mini-Tn) donor plasmid (Fig. 1b). Targeted insertions are generated in a single step through conjugative delivery of both plasmids into a recipient bacterial strain. Amplification and sequencing of the inserted guide sequence is then used as a proxy for mutant abundance.

During the development of MultiCAST, we identified several factors that influence the observed CAST insertion frequency in pooled mutant populations and devised strategies to mitigate some of their effects. First, we found that guide-transposon “crosstalk” occurs when a guide RNA expressed from one transposon directs insertion of a different guide-containing transposon present on another plasmid within the same cell, leading to an overestimation of low-frequency insertions by guide-based amplicon sequencing. We resolved this problem by diluting the mini-Tn donor strain prior to conjugation, which reduces the likelihood of multiple plasmids entering into the same recipient cell. Second, we discovered that chromosomal regions bound by H-NS exhibit reduced insertion frequencies, indicating that H-NS physically blocks Tn6677 transposition. Third, we observed substantial variability in guide activity and developed a machine learning model to distinguish highly active guides from low activity ones based on target sequence features. Together, our findings provide fundamental insights into the behavior of this exciting class of mobile genetic elements. These insights have directly informed the design of MultiCAST, resulting in a robust genetic screening platform. We anticipate that they will also guide the development of future CAST systems for bacterial genome engineering and functional genetics.

## Results

### A conditionally replicative CAST system for single-step transposon insertion

The Type I-F3 CRISPR-associated transposon (CAST) Tn6677 encodes three CRISPR-Cas proteins (Cas8, Cas7, and Cas6) that together form the guide RNA-directed DNA targeting complex known as Cascade (27). Upon recognizing a DNA target site (protospacer) adjacent to a 5’-CN-3’ protospacer adjacent motif (PAM), where N represents any nucleotide, Cascade recruits the Tn7-like transposition machinery (TnsA, TnsB, and TnsC) to the site via the bridge protein TniQ (28). Once recruited, the transposon, flanked by defined right (R) and left (L) end sequences, is inserted ~49 bp downstream of the 3’ end of the target site by TnsB (Fig. 1a).

Tn6677 was initially adapted by Vo et. al. (7) into a single-plasmid system for targeted transposition termed pSPIN, in which all necessary components, including the CRISPR-Cas proteins, transposition proteins, guide RNA, and transposon DNA, are encoded on a single, constitutively replicative plasmid that can be introduced into a recipient bacterial strain by transformation or conjugation. We adapted pSPIN into a conditionally replicative, two-plasmid system called MultiCAST (Fig. 1b). In this configuration, all seven proteins required for targeted transposition (TnsABC, TniQ, and Cas876) are encoded on a separate “helper” plasmid, while a mini-transposon (mini-Tn), flanked by R and L sequences and containing an antibiotic selectable marker, is encoded on a mini-Tn donor plasmid. The mini-Tn contains a cloning site for Gibson assembly of a ssDNA oligonucleotide encoding the guide sequence (29).

Due to their R6K origin, both plasmids used in MultiCAST are non-replicative in most bacterial strains, which lack the π protein required for propagation (30). A conditionally replicative two-plasmid system offers several advantages over the original design by Vo et. al. and related systems that rely on a single plasmid or constitutively replicative origins. Delivery of the CAST machinery and transposon on conditional replicons ensures that all transconjugants (defined as antibiotic-resistant colonies arising post-conjugation) are derived from transposon insertion mutants. In contrast, systems using constitutively replicative plasmids require plasmid curing after transposition to confirm that transconjugants represent transposon insertions in the genome. Without curing, low-throughput methods such as quantitative PCR must be used to determine whether the transposon has successfully inserted into the target locus. Additionally, separating the guide-containing transposon from the enzymatic machinery prevents self-targeting in the donor strain, which can lead to guide dropout. Thus, the use of a conditionally replicative two-plasmid system enables direct selection for transposon insertions, eliminating the need for post-insertion plasmid curing, and prevents the loss of guides targeting essential genes. The design of MultiCAST is similar to the two-plasmid system developed by Banta et. al. (21).

After guide oligo assembly into the mini-Tn donor plasmid and transformation into a Pir^+^ donor strain, both plasmids are introduced into the recipient strain via conjugation. We use the Pir^+^ *E. coli* donor strain MFD*pir*, a diaminopimelic acid (DAP) auxotroph, which allows for efficient counterselection of the donor strain by omitting DAP from the medium (31). Consequently, colonies that form on selective medium following conjugation represent recipient cells harboring the transposon inserted into the genome.

### Single-gene disruption using MultiCAST

We first validated MultiCAST as a method for single-gene inactivation using a guide targeting the *lacZ* gene in *E. coli* MG1655. Disruption of *lacZ*, which encodes β-galactosidase, results in white colonies instead of blue using growth medium containing 5-bromo-4-chloro-3-indolyl-β-D-galactopyranoside (X-gal). Conjugation for 2 h at 37 °C followed by immediate plating on selective medium yielded an average transposition frequency of 2.8×10^−4^ (defined as transconjugants per total recipient cells; Fig. 1c). Given prior observations that lower temperatures enhance CAST activity (7), we tested the effects of post-conjugation recovery time and temperature on transposition frequency. Extended recovery at room temperature progressively increased transposition frequency by ~1,000-fold, reaching ~3.1×10^−1^ after 25 h. In contrast, extended recovery at 37 °C did not increase the number of mutants (Fig. 1c). Importantly, no colonies emerged when either the donor or helper plasmid was omitted during the conjugation, confirming that colony formation was entirely CAST-dependent (Fig. 1d).

Phenotypic screening of transconjugants on X-gal plates revealed 100% *lacZ* inactivation (15/15 colonies; Fig. 1d). Orientation-specific PCR across 48 colonies demonstrated exclusive insertion of the transposon in the ‘RL’ orientation (right end 5’ to left end; Fig. 1f). This strong directional bias has been previously reported for Tn6677 (7), and we anticipate that the RL orientation will generally predominate across target sites.

Based on these data, we conclude that MultiCAST is an efficient and specific method for single-gene disruption. Two of its key advantages are speed and simplicity: mutants can be generated in as little as three days from donor plasmid assembly to colony isolation, with minimal hands-on time. We next sought to evaluate MultiCAST for pooled gene disruption.

### Pooled gene disruption using MultiCAST

To evaluate MultiCAST for pooled gene inactivation, we assembled a pool of 5,225 guide oligos targeting 88 genes in *E. coli* MG1655 into the mini-Tn donor plasmid (Fig. 2a, Dataset S1). Guides were designed to target both the template and non-template strands within the first half of each gene to maximize the likelihood of functional disruption. The only other constraint was the presence of a 5’-CN-3’ PAM upstream of the target sequence. Amplicon sequencing of the cloned guides revealed that all but two were represented by at least one read in the donor pool, while 5,190 (99.3%) were represented by at least 100 reads (Fig. 2b).

**Fig. 2.**
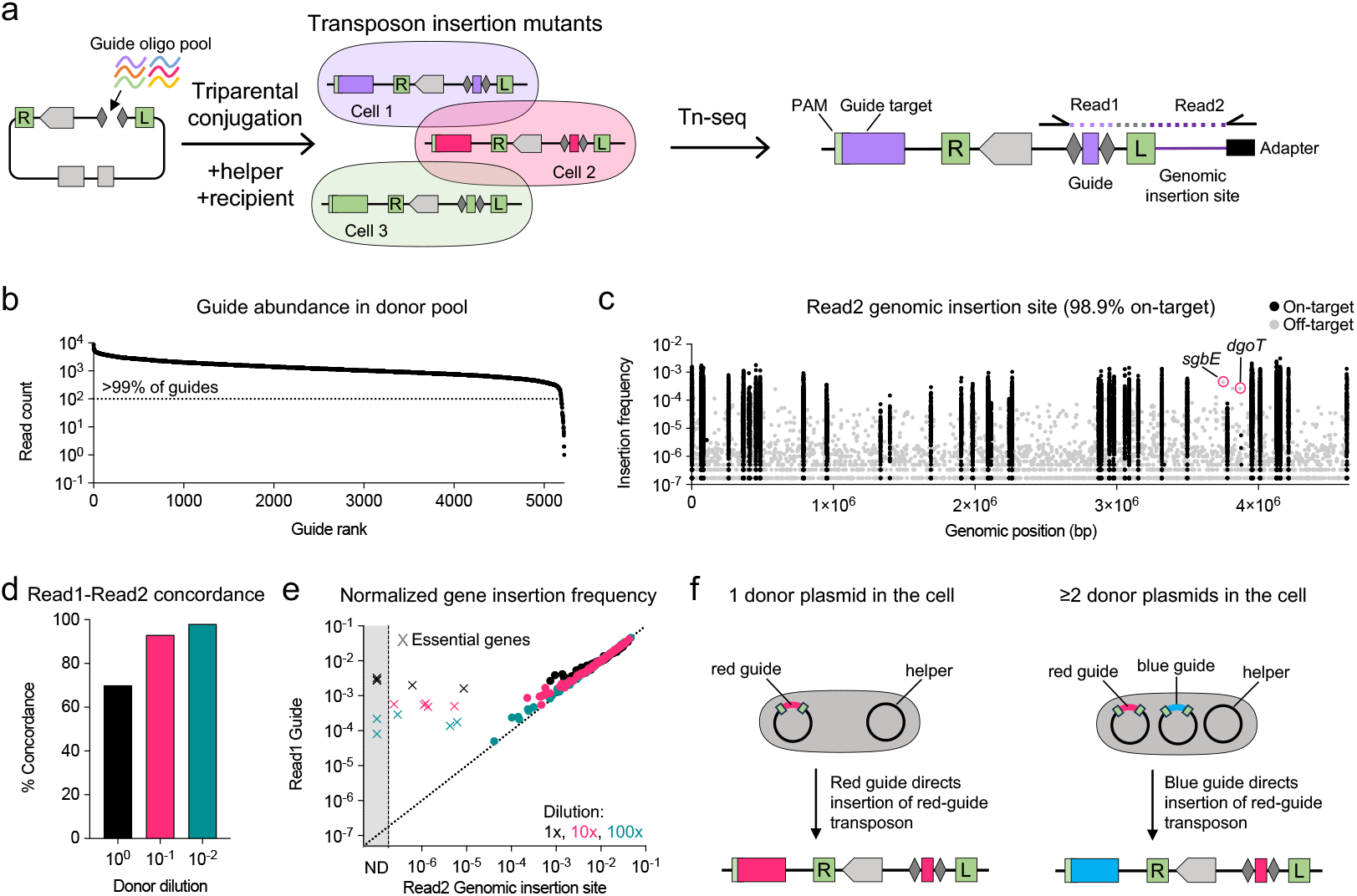
Pooled gene knockout using MultiCAST. **a)** Illustration of the pooled targeting and sequencing approach. A pool of guide oligos is assembled into the linearized mini-Tn donor plasmid. Following conjugation and plating on selective medium, the recipient population contains diverse CAST-mediated insertions, which are quantified by Illumina paired-end sequencing following Tn-seq library preparation. Read1 captures the guide sequence while Read2 captures the genomic insertion site. **b)** Distribution of guide representation in the donor pool. Of the 5,225 designed guides, 99.3% (5,190) were detected at >100 reads. **c)** CAST insertion frequencies across the *E. coli* MG1655 genome. Each dot represents a unique insertion site. Black dots indicate on-target insertions within the 88 targeted genes; gray dots indicate off-target insertions outside these genes. Circled gray dots denote the two most frequent off-target insertion sites, likely due to partial guide matching to the off-target site. **d)** Concordance between Read1 and Read2 at different dilutions of the mini-Tn donor pool prior to conjugation. Concordant reads are defined as Read2 reads mapping to the gene targeted by the guide in Read1. Dilution improves concordance by reducing donor plasmid crosstalk. **e)** Comparison of normalized gene insertion frequencies derived from Read1 and Read2 at different donor pool dilutions. Each dot represents a targeted gene. For each gene, read counts were summed for all guides targeting the gene (Read1) or inserted at the gene (Read2) and normalized to both the total number of reads in the sample and the number of guides targeting that gene. **f)** Illustration of donor plasmid crosstalk. See main text for details.

To assess whether amplicon sequencing of the guide could serve as a proxy for mutant abundance, we compared guide frequency to the ground truth transposon insertion frequency determined by Tn-seq, which identifies the exact junction between the transposon and the genome. Following selective plating of transconjugants on lysogeny broth (LB) agar plates, genomic DNA was extracted and prepared for Illumina paired-end sequencing using a modified Tn-seq library preparation protocol (Methods). In this approach, Read1 captures the cloned guide sequence within the transposon, while Read2 identifies the genomic insertion site. This design enables a direct comparison between the frequency of each guide in Read1 and the corresponding genomic insertion site in Read2 (Fig. 2a).

Analysis of Read2 genomic insertion sites revealed that 98.9% of mapped reads were on-target, defined as insertions within one of the 88 targeted genes (Fig. 2c). This high on-target rate is consistent with the well-characterized specificity of Tn6677-mediated transposition (7, 21). Among off-target events, the most frequent insertion occurred within the gene *sgbE*, which shares 75% nucleotide identity with *araD*, one of the targeted genes (Fig. 2c, Fig. S1a). A single guide targeting *araD* accounted for nearly all insertions into *sgbE*, with 25 of 32 positions matching a putative target site in *sgbE* (Fig. S1a). Surprisingly, the off-target site is preceded by ‘GC’ rather than the canonical ‘CN’ PAM. Of the seven mismatches, four occurred at the sixth nucleotide positions of the guide sequence, a position known to be flipped out during Cascade binding and thus dispensable for target recognition by Type I-F CAST systems (28, 32). ~23% of Read1 reads containing this specific *araD* guide were inserted at the *sgbE* locus.

The second most frequent off-target insertion occurred within the gene *dgoT* (Fig. 2c). A single guide targeting *araA* accounted for nearly all insertions in this case. However, the associated guide sequence from Read1 exhibited only 21 of 32 matches to a putative target site, with several mismatches occurring outside the sixth positions (Fig. S1b). Together, these findings underscore that while Tn6677 exhibits high specificity relative to other CAST systems, it retains some flexibility in its target recognition (32). This tolerance for mismatches and noncanonical PAMs may provide a selective advantage in the spread of this mobile genetic element. However, rare off-target transposition observed with certain guides could lead to inaccurate estimates of mutant abundance.

### Guide-transposon crosstalk overestimates mutant abundance

Comparison between the Read1 guide sequence and the Read2 insertion site revealed ~70% concordance (Fig. 2d, black bar). When individual guide frequencies from Read1 were aggregated per gene and compared to the corresponding insertion frequencies from Read2, we observed that the guide sequence overestimated the abundance of mutants compared to Tn-seq, particularly for essential genes (Fig. 2e).

We hypothesized that discordance between Read1 and Read2, along with the overestimation of mutant abundance by the guide in Read1, arises from plasmid “crosstalk” in which a single recipient cell receives multiple mini-Tn donor plasmids containing distinct guide sequences during conjugation (Fig. 2f). In such cases, any mini-Tn can be excised by the transposase, regardless of which guide is encoded within the transposon. Consequently, the guide RNA expressed from one donor plasmid may direct insertion of a mini-Tn from another donor plasmid, resulting in mismatched guide-insertion pairs and artificially inflated guide frequencies for low-abundance mutants.

To mitigate crosstalk, we diluted the donor strain carrying the guide pool prior to conjugation, reasoning that this would lower the likelihood of multiple mini-Tn donor plasmids entering into the same recipient cell. 10x and 100x dilutions improved both the concordance between Read1 and Read2 (Fig. 2d) and the correlation between guide frequency and insertion frequency (Fig. 2e). To further confirm the impact of crosstalk, we performed an orthogonal experiment by mixing a *lacZ*-targeting donor strain 1:1 with a non-targeting donor (empty vector) and serially diluting the mixture prior to conjugation. PCR-based sequencing of the resulting transconjugant population revealed that non-targeting donor reads decreased from 17% (no dilution) to just 0.09% at 1,000-fold dilution (Fig. S2).

Despite this improvement, guide frequency continued to overestimate mutant abundance for essential genes, likely due to residual crosstalk as well as off-target insertions. In many cases, these off-target events were not driven by partial guide-target complementarity, but rather by an unusually long distance between the guide-targeted site and the actual insertion site, resulting in transposon insertion outside the coding sequence of the essential gene (Fig. S1c).

Collectively, these results demonstrate that guide frequency can serve as a reliable proxy for mutant abundance when crosstalk is minimized. However, for essential genes, guide-based measurements alone may significantly overestimate mutant abundance, necessitating direct measurement of CAST genomic insertion sites with Tn-seq for accurate quantification.

### H-NS inhibits CAST insertion

We observed substantial variability in CAST insertion frequency across non-essential genes, spanning over three orders of magnitude (Fig. 3a). For example, *lacA* and *lacY*, despite sharing an operon with *lacZ*, exhibited >100-fold lower insertion frequencies. Similarly, *cas3*, the non-essential Type I-E CRISPR-Cas nuclease, was among the least frequently inserted genes in the pool (Fig. 3a).

**Fig. 3.**
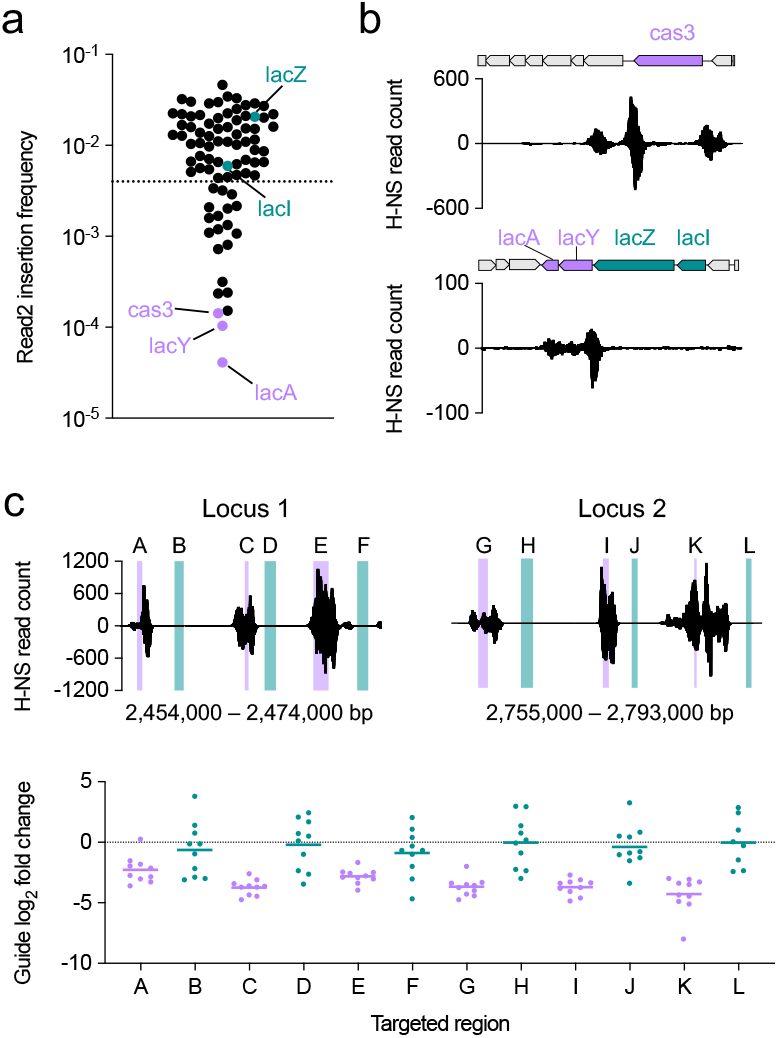
H-NS inhibits CAST insertion. **a)** Gene-level insertion frequencies quantified from Read2 data at 100x donor dilution. Guides targeting genes above the dotted line were used to train a machine learning model. **b)** Locus-specific H-NS binding profiles. Positive and negative values represent read counts mapped to the plus and minus DNA strands, respectively. **c)** Guide activity at H-NS-bound regions compared to adjacent unbound regions. Activity is expressed as the log_2_ fold change in guide frequency between the recipient and donor populations. The horizontal line indicates the mean guide activity within each targeted region. H-NS binding data were obtained from Choe et. al. (25).

The *cas3* gene in *E. coli* K12 is transcriptionally repressed by the nucleoid-associated protein H-NS (33). To investigate whether H-NS binding was associated with the reduced insertion frequencies, we examined chromatin immunoprecipitation (ChIP-Exo) data from Choe et. al. (25), which revealed high H-NS occupancy at both the *cas3* locus and the lac operon (Fig. 3b). Notably, H-NS binding specifically overlapped *lacA* and *lacY* while sparing *lacZ*.

H-NS is known to inhibit various classes of transposons (24, 25), and we hypothesized that it may also inhibit CAST-mediated transposition. A direct test of this hypothesis would be to measure CAST activity at H-NS-bound loci in an *hns* mutant background. However, functional redundancy between nucleoid-associated proteins, particularly between H-NS and its paralog StpA, could confound the result (25). Moreover, disruption of *hns* causes widespread phenotypic effects, potentially confounding the measurement of conjugation and transposition efficiency (34).

To circumvent these challenges, we used an orthogonal approach to assess the causal role of H-NS in inhibiting CAST activity. We designed a pool of guides targeting two non-essential chromosomal loci in *E. coli* MG1655 (35), each containing three regions of high and low H-NS binding (Fig. 3c). Ten guides were designed to target each region, yielding a total of 120 guides (60 per locus; Dataset S2). Because H-NS preferentially binds AT-rich DNA (36), we minimized differences in GC content between the two groups to mitigate guide sequence composition as a confounding variable (48.5% GC in bound regions vs. 49.8% in unbound regions; Fig. S3).

Transposition frequency was quantified by amplicon sequencing of the guide, and activity was defined as the log_2_ fold change of guide frequency in the recipient population divided by the donor (Fig. 3c). As predicted, guides targeting H-NS-bound regions exhibited consistently lower average activity than those targeting adjacent unbound regions across both loci, demonstrating that H-NS directly inhibits CAST insertion. These findings reveal that CAST insertion efficiency is modulated by DNA-binding proteins such as H-NS and underscore the importance of distinguishing insertion efficiency from mutant fitness when interpreting phenotypic data (25). Notably, transcription is another factor influencing guide activity, potentially due to collisions between the CAST machinery and RNA polymerase (21).

### DNA sequence features predict CAST activity

We observed substantial variability in the insertion frequencies of individual guides targeting the same gene, with log_2_ fold changes spanning over three orders of magnitude between the least and most active guides (Fig. 4a). This variability occurred for guides targeting both the template and non-template strands, with no consistent strand-specific difference in average activity across genes. In some cases, even overlapping guides exhibited markedly different activities, suggested that guide activity is not solely dictated by physical barriers such as DNA-binding proteins or transcriptional interference (21).

**Fig. 4.**
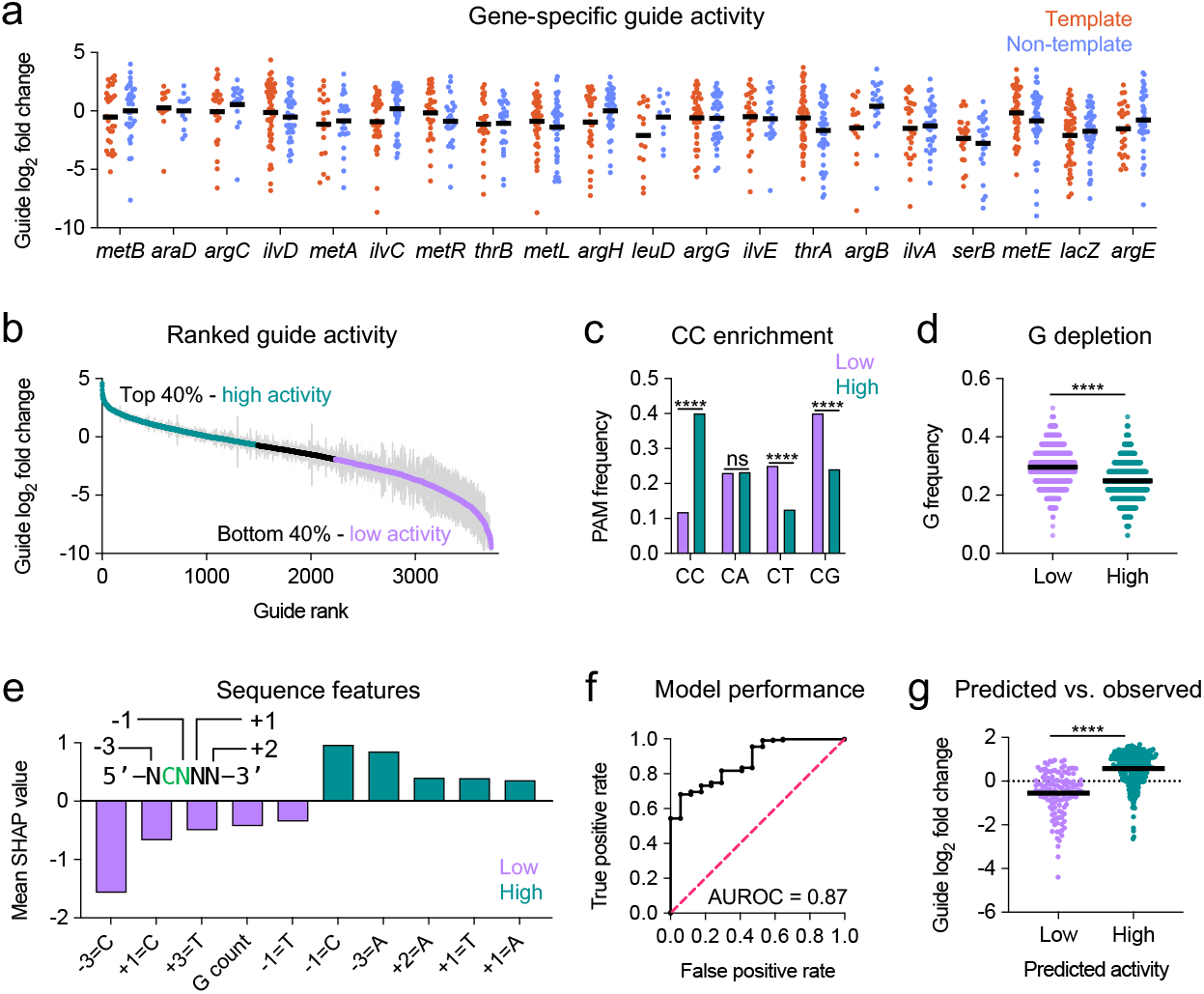
DNA sequence features predict guide activity. **a)** Guide activity for the top 20 genes with the most abundant insertions from Fig. 3a, separated by the targeted strand. **b)** Ranked guide activity for all guides with ≥100 reads in the donor pool, targeting genes above the dotted line in Fig. 3a. Guides were classified into high-activity (top 40%) and low-activity (bottom 40%) groups and used to train the machine learning model. Each point represents the mean ± SEM of three biological replicates. **c)** PAM sequence frequencies across high- and low-activity guide groups. Asterisks indicate *P* <0.0001 by Fisher’s exact test; ns = not significant. **d)** Guanine nucleotide frequency is reduced in high-activity guides. The horizontal line indicates the mean. Asterisks indicate *P* <0.0001 by two-sided Mann-Whitney test. **e)** SHAP analysis showing the top five sequence features associated with high and low guide activity. The first nucleotide of the guide is designated as +1. **f)** External validation of the predictive model using a pool of 385 guides. AUROC = area under the receiver operating characteristic curve. **g)** Predicted vs observed guide activity in the external validation dataset. The horizontal line indicates the mean. Asterisks indicate *P* <0.0001 by two-sided Mann-Whitney test.

We hypothesized that DNA sequence features at the target site might influence CAST activity. To identify these features, we ranked average guide activities for genes with insertion frequencies >4×10^−3^ (62 out of 88 genes; all genes above the dotted line in Fig. 3a). Genes with higher insertion frequencies were selected to minimize confounding effects from DNA-binding proteins and fitness-related factors. Guides were then classified into high activity (top 40%) and low activity (bottom 40%) groups, each containing 1,492 guides (Fig. 4b). In addition to the 32 bp guide sequence, we examined the 5 bp upstream of each guide target site, including the PAM (positions −1 and −2) and adjacent upstream bases (positions −3 to −5).

One of the most striking differences between high- and low-activity guides was PAM composition. High-activity guides were significantly enriched for a CC PAM compared to low-activity guides (40.0% vs. 11.9%, *P* < 0.0001). In contrast, CT and CG PAMs were more prevalent among low-activity guides (25.1% and 40.0% vs. 12.6% and 24.1%, respectively; *P* < 0.0001), while CA PAM frequencies did not differ significantly between groups (Fig. 4c). Additionally, high-activity guides exhibited a lower overall guanine (G) content within the guide sequence (24.9% vs. 29.6%, *P* < 0.0001; Fig. 4d), a bias that may relate to stability of R-loop formation (37).

To systematically identify sequence features predictive of CAST activity, we trained an Extreme Gradient Boosting (XGBoost) classifier (38) using 239 sequence-derived features to distinguish high-from low-activity guides (Methods). The model demonstrated strong predictive performance on the training dataset, achieving a mean area under the receiver operating characteristic curve (AUROC) of 0.89 in 10-fold cross-validation and 0.91 in leave-one-gene-out (LOGO) validation (Fig. S4a and b; Tables S1 and S2), indicating robust generalization to unseen genes.

We performed SHAP (SHapley Additive exPlanations) analysis to quantify feature importance (39). Consistent with our earlier findings, the presence of a cytosine at position −1 (forming a CC PAM, given the invariant C at position −2) was among the strongest positive predictors of activity (Fig. 4e). Conversely, thymine at position −1 and higher G content within the 32-nt guide was negatively associated with activity. SHAP analysis also revealed additional position-specific preferences: adenine at positions −3, +1, and +2, and thymine at +1, correlated with high activity, while thymine at +3, and cytosine at −3 and +1, correlated with low activity (Fig. 4e, Fig. S4c, Dataset S3). Interestingly, cytosines immediately adjacent to the PAM (positions −3 and +1) were strongly predictive of low activity (Fig. 4e). We speculate that the presence of a C nucleotide at these positions may interfere with CRISPR-Cas recognition of the PAM, potentially causing “slipping” of the complex and impaired targeting.

To independently validate our machine learning model, we tested a new set of 385 guides targeting two genes inserted at the chromosomal *att*Tn7 site (40) that were not included in the training data (Dataset S4). The model achieved an AUROC of 0.87 (Fig. 4f), and guides predicted to be highly active exhibited significantly higher measured activity (Fig. 4g, *P* < 0.0001; Table S3). Moreover, predicted probabilities correlated with observed activity across all guides (Spearman’s ρ = 0.682, *P* < 0.0001), indicating that the model effectively captures activity rankings to further inform guide selection (Fig. S4d). A web portal implementing the trained model is available at https://multicastguidepredictor-v1.streamlit.app and can be used to choose guide sequences for genes of interest.

Together, these findings demonstrate that CAST activity is primarily governed by sequence features immediately upstream and downstream of the PAM, with the guanine content within the guide sequence also contributing to activity.

### High-throughput genetic screening with MultiCAST

To evaluate MultiCAST as a platform for high-throughput genetic screening, we assayed our 88-gene mutant population across twelve nutrient conditions and measured fitness using PCR-based guide sequencing. Three independently generated mutant populations were tested per condition, yielding a total of 36 samples (Fig. 5a). Condition-specific fitness effects were readily apparent. Mutants in genes essential for amino acid biosynthesis were depleted in minimal medium, with pathway-specific rescue upon supplementation with the corresponding amino acid (Fig. 5b). Similarly, mutants in carbon utilization pathways were selectively depleted when grown with their respective substrates as the sole carbon source, while mutants in pathway repressors were enriched. These results validate MultiCAST as a robust platform for efficient, high-throughput genetic screening, enabling systematic interrogation of gene function at scale.

**Fig. 5.**
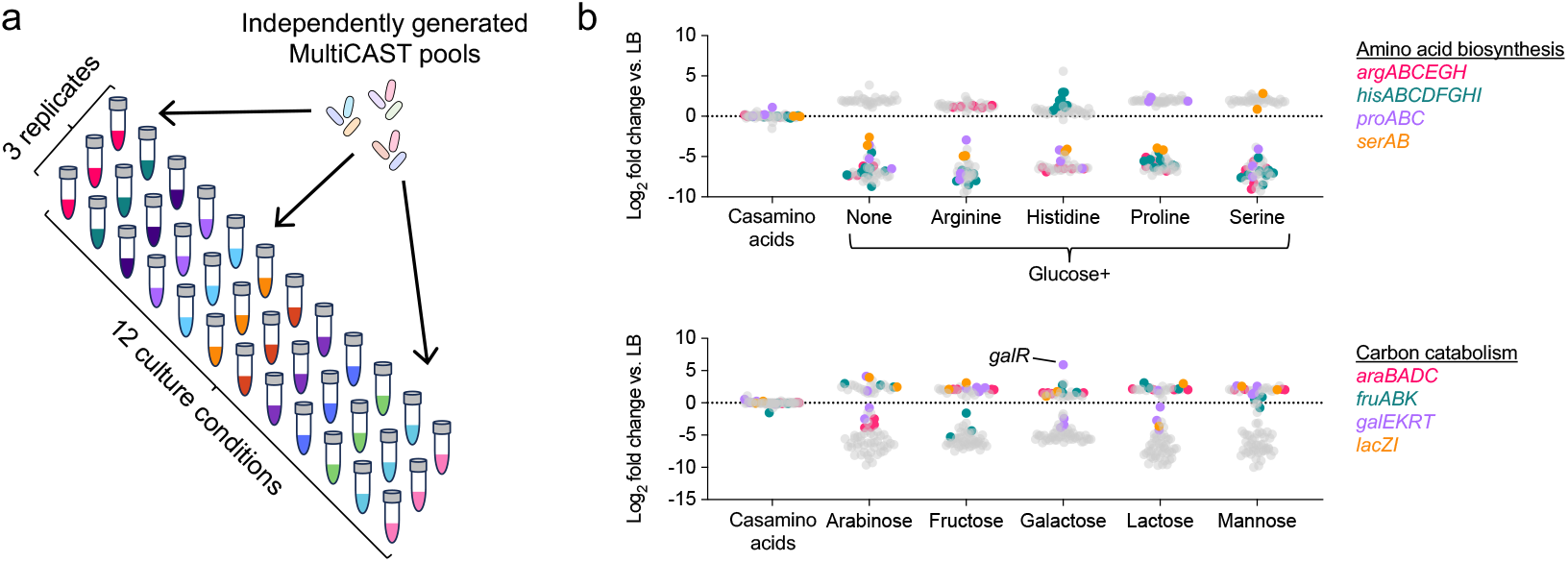
High-throughput genetic screening with MultiCAST. **a)** Diagram of the screen to test gene requirements across twelve nutrient conditions. Three independently generated CAST mutant populations were assayed in each condition for a total of 36 samples. Insertions were quantified by amplicon sequencing of the guide. **b)** Results of the screen showing the average log_2_ fold change in insertion frequency for each gene in the specified nutrient condition relative to LB. Colored dots indicate genes previously reported to affect fitness in the specified medium (53). *galR* encodes a repressor of galactose utilization.

## Discussion

CRISPR-associated transposons hold great promise for genome engineering and functional genetics in both bacterial and eukaryotic organisms (41, 42). In this study, we adapted the Type I-F3 CAST Tn6677 into a streamlined platform called MultiCAST, enabling single-step transposon mutagenesis to generate individual or pooled targeted mutants in just a few days. By incorporating the guide sequence directly into the mini-Tn donor, MultiCAST creates an easily readable molecular tag that enables straightforward quantification of mutant frequency via PCR-based amplicon sequencing. We benchmarked MultiCAST against Tn-seq, the gold standard for transposon insertion profiling, and observed strong concordance for non-essential genes. In contrast to Tn-seq, the simplicity of generating sequencing libraries using a single PCR reaction makes MultiCAST well-suited for high-throughput functional screens.

While transposon insertion frequency is often interpreted as a proxy for mutant fitness, our study highlights several fitness-independent factors that influence CAST activity. We found that the nucleoid-associated protein H-NS inhibits transposition, consistent with its known role in suppressing other transposon systems (24, 25). Prior work has also shown that host factors such as IHF and RecD also modulate CAST activity (43, 44), and has implicated RNA polymerase collisions at highly transcribed loci as a barrier to CAST transposition (21). It is possible that other DNA-binding proteins, including transcriptional regulators, may similarly affect CAST activity.

In addition to DNA binding proteins, sequence features at the target site significantly impact CAST activity. We systematically identified sequence features associated with high and low guide activity, finding that the identity of the PAM, as well as the immediate upstream and downstream sequences, strongly influence activity. Guanine content of the guide sequence also correlates with activity. Together, these findings underscore the importance of accounting for non-fitness-related factors when designing and interpreting functional genetic screens.

We encountered two key challenges in accurately quantifying transposon insertion frequency by guide sequencing: plasmid crosstalk, where multiple mini-Tn donor plasmids enter the same recipient cell, and off-target insertion driven by partial guide-target complementarity. We found that crosstalk could be mitigated by diluting the donor strain prior to conjugation, which improved accuracy but proportionally reduced transposition efficiency. Although the overall off-target rate was low (~1%), it could still confound interpretation for guides targeting loci with low on-target efficiency due to DNA-binding protein barriers such as H-NS.

In future incarnations of MultiCAST, it should be possible to circumvent both crosstalk and off-target insertion by incorporating random barcodes into the mini-Tn donor plasmid. These barcodes can then be linked to genomic insertion sites via Tn-seq. This approach, pioneered by Wetmore et. al. in a method called RB-Tn-seq (19), would allow PCR-based quantification of mutant frequency using the barcode sequence rather than the guide sequence. Combining RB-Tn-seq with MultiCAST would leverage the strengths of both methods: accurate mutant identification using barcodes linked to the ground truth insertion site, and targeted and compact library design using CRISPR RNA guides. The relatively small size mutant populations generated using a targeted approach, as opposed to random transposition, are particularly advantageous for genome-scale screening in bottlenecked experimental systems (35).

We used the Type I-F3 CAST Tn6677 from *Vibrio cholerae* strain HE-45, one of the earliest described CAST systems (4). Type I-F3 systems are attractive genetic tools due to their high specificity, but properties such as PAM preference, insertion site distance, and orientation bias vary widely even within this subtype (45). For example, Tn7479 from *Psychromonas* spp. exhibits near PAM-less targeting and an exceptionally strong RL orientation bias, making it ideal for applications requiring directional insertion, such as in-frame protein tagging (43, 45). Our study provides a framework for evaluating alternative CAST systems for high-throughput genetic screens. In parallel, engineering or directed evolution of CASTs to enhance specificity and activity represents a promising avenue for future development (13, 26, 46), similar to the efforts that have transformed CRISPR-Cas9 into a powerful genome editing tool (47).

Given the demonstrated efficacy of CAST systems across diverse bacterial species, we anticipate that MultiCAST and related platforms will become standard tools for bacterial genetics and genome engineering. By eliminating experimental barriers to mutant creation and sequencing library preparation, MultiCAST offers a reliable and accessible method for genome-scale functional screening, expanding the scope and scalability of genotype-phenotype studies in microbiology.

## Materials and Methods

### Plasmid assembly

All plasmids were assembled with the Gibson method using the NEBuilder HiFi DNA Assembly Master Mix (New England Biolabs, #E2621)(29). DNA fragments were amplified using primers from Integrated DNA Technologies (IDT). The CAST enzymatic machinery from *Vibrio cholerae* was amplified from pSL1765 (Addgene #160734) (7). Assembled plasmids were electroporated into *E. coli* strain MFD*pir* (31) and purified using the Monarch Spin Plasmid Mini Prep Kit (New England Biolabs, #T1110). Plasmid sequences were verified by whole-plasmid sequencing (Plasmidsaurus) and edited using ApE (A Plasmid Editor) (48). The MultiCAST mini-Tn donor and helper plasmid sequences will be made available on Addgene.

### Guide oligo assembly into mini-Tn donor

Purified mini-Tn donor plasmids were digested with PacI and SpeI (New England Biolabs, #R0547 and #R3133). Guide oligos were assembled into the linearized plasmid using NEBuilder HiFi DNA Assembly Master Mix at a 5:1 molar ratio (oligo:plasmid), then electroporated into E. coli MFD*pir*. For large oligo pools, initial cloning into a high-competence Pir^+^ strain is recommended before transformation into MFD*pir*. Following transformation, cells were incubated at 37 °C for 1 h in 1 ml LB+30 μM diaminopimelic acid (DAP), then plated on LB+DAP+50 μg/ml kanamycin and grown overnight with shaking at 37 °C.

For single-gene knockout, single-stranded DNA oligos from IDT were used directly in the Gibson assembly. For pooled knockouts, oligos from Twist Biosciences were amplified to generate sufficient quantities for assembly. Each oligo contains a 32-nt guide sequence flanked by homology arms for assembly into the linearized donor plasmid:

5’-<u>TACTACTGCAAAGTAGCTGATAAC</u>-**[32 nt guide]**-<u>CTTTACTGCTGAATAAGTAGATAACTAC</u>-3’

The underlined regions correspond to atypical CRISPR repeats flanking the guide RNA (32, 49).

### Conjugative delivery of MultiCAST plasmids

Overnight cultures of MFD*pir* strains carrying the mini-Tn donor plasmid (with cloned guide), the helper plasmid, and the recipient strain were pelleted 2 min at 8,000 g, washed once with LB, and resuspended in 100 μl LB. Equal volumes of each strain were mixed 1:1:1. For pooled knockouts, the donor strain was diluted 100x prior to mixing to reduce plasmid crosstalk.

A 60 μl mating mixture was spotted onto a 0.45 μm mixed cellulose ester filter disc (Millipore #HAWP02400) placed on an LB+DAP plate. After 2 h incubation at 37 °C, the filter was transferred to 1 ml LB in a microfuge tube and vortexed to resuspend cells. The resuspension was incubated at room temperature with gentle agitation from 4 h to overnight (longer incubation improves transposition frequency). Following recovery, cells were plated on LB+50 μg/ml kanamycin. Resulting colonies represent targeted transconjugants.

### Transposon insertion sequencing

Paired-end Tn-seq libraries were prepared as described previously (35), with minor modifications. Genomic DNA was extracted using the DNeasy Blood and Tissue Kit (Qiagen #69504), then sheared to ~500 bp using an M220 ultrasonicator (Covaris). Sheared DNA was end-repaired using the Quick Blunting Kit (NEB #E1201), A-tailed with Taq polymerase (NEB #M0273), and ligated to hybridized Illumina P7 adapters with complementary T-overhangs using T4 ligase (NEB #M0202).

200 ng of ligated DNA was amplified for 20 cycles using a transposon-specific forward primer and an adapter-specific reverse primer containing a unique i7 index. Libraries were size-selected using SPRIselect beads (Beckman Coulter #B23318) at a 0.65x ratio, quantified with a Qubit fluorimeter, pooled, and sequenced on an Illumina Nextseq 1000/2000 using a 300-cycle kit. Read1 (65 cycles) captured the guide sequence while Read2 (235 cycles) captured the transposon insertion site. Primer sequences for Tn-seq are listed in Table S4.

### High-throughput phenotypic screening

Frozen 50 μl aliquots of three replicate 88-gene MultiCAST mutant pools in *E. coli* MG1655 were thawed and diluted into 5 ml LB+50 μg/ml kanamycin, then grown overnight at 37 °C. Cultures were washed twice with 1 ml phosphate buffered saline (PBS), resuspended in 1 ml PBS, diluted 1:1000, and 100 μl was used to seed 3 ml of fresh LB or M9 medium containing 0.2% of the indicated carbon sources, with or without 0.01% of the indicated amino acids. Cultures were grown overnight at 37 °C.

### Guide amplicon sequencing

Guide sequencing libraries were prepared as previously described (50). Mutant populations were diluted in water, and guide sequences were amplified using 2 μl of diluted cells and One*Taq* 2x Master Mix (NEB #M0482) with the following thermocycler protocol:

1. 94°C for 30 s
2. 94°C for 20 s
3. 60°C for 30 s
4. 68°C for 30 s (Repeat steps 2 - 4 for 25 cycles)
5. 68°C for 5 min

Primer sequences for pooled guide amplification are listed in Table S4. All primers share identical binding sites flanking the guide sequence but contain unique indices to enable sample multiplexing. The resulting amplicons range from 261 to 268 bp in length, including the sequencing adapters added during PCR. The slight variation in amplicon size is due to the variable region within each indexing (var) primer, which introduces sequence diversity for improved sequencing performance. PCR products were pooled and purified using QIAquick reagents (Qiagen), quantified with a Qubit fluorimeter, and sequenced on an Illumina NextSeq 1000/2000 using 100-cycle single-end sequencing.

### Machine learning model for classifying guide activity

We trained an Extreme Gradient Boosting (XGBoost) classifier (38) to distinguish high-from low-activity guides. Guide labels were assigned based on the activity distribution in the MultiCAST Pool1 dataset: the top 40% were labeled as high-activity (positive class), and the bottom 40% as low-activity (negative class). Guides in the middle 20% were excluded from training to establish a clear decision boundary for the model.

Each guide was encoded using two feature classes: (i) positional one-hot encoding of each nucleotide at every position within the PAM region and the 32-nt guide sequence, and (ii) global sequence features including GC content, per-base counts (A/C/G/T), homopolymer run lengths for each base, guide-level Shannon entropy, per-base counts within the PAM region; and exact k-mer counts for k=2 and k=3. In total, 239 features were used for model training.

Internal validation was performed using 10-fold cross-validation, where the dataset was randomly partitioned into ten subsets, and leave-one-gene-out (LOGO) validation, where all guides targeting a single gene were held out for testing in each fold. For each validation strategy, we reported the mean ± S.D. of the following metrics: area under the receiver operating characteristic curve (AUROC), area under the precision-recall curve (AUPRC), precision, recall, and F1 score (Tables S1 and S2).

Model selection was performed using an inner 5-fold StratifiedKFold (52) randomized search over XGBoost hyperparameters (n_estimators, learning_rate, max_depth, min_child_weight, subsample, colsample_bytree, reg_alpha, reg_lambda, gamma, max_delta_step, and grow_policy). The F1 score was used as the selection metric.

To generate the final model for prospective guide prediction, we re-tuned hyperparameters using GroupKFold (52) by gene, selecting the configuration with the highest mean cross-validated F1 score across held-out genes. The final model was evaluated on an external dataset comprising 385 guides (Dataset S4 and Table S3).

To interpret feature contributions, we computed Shapley Additive Explanations (SHAP) values using shap.TreeExplainer (39). For each feature, importance was summarized by mean absolute SHAP value, mean positive and mean negative SHAP values, and Spearman correlation between the feature value and its SHAP value. The Spearman correlation provides the direction of association between the feature and its contribution to the positive class. We reported the top five positively associated features as those with Spearman correlation >0 and the highest mean positive SHAP values, and the top five negatively associated features as those with Spearman correlation <0 and the lowest mean negative SHAP values.

All analyses were performed in Python (v3.13.7) using the scikit-learn (52), shap (39) and xgboost (38) modules. Code and custom scripts are available at https://github.com/YiyanYang0728/MultiCAST_scripts and https://github.com/YiyanYang0728/MultiCAST_guide_predictor.

### Chromosomal attTn7 site insertion

Overnight cultures of MFD*pir* strains carrying a mini-Tn7 donor plasmid containing the chloramphenicol and kanamycin resistance genes, the helper plasmid pJMP1039 (Addgene plasmid #119239) (51), and the recipient strain MG1655 were pelleted 2 min at 8,000 g, washed once with LB, and resuspended in 100 μl LB. Equal volumes of each strain were mixed 1:1:1.

A 60 μl mating mixture was spotted onto a 0.45 μm mixed cellulose ester filter disc (Millipore #HAWP02400) placed on an LB+DAP plate. After 2 h incubation at 37 °C, the filter was transferred to 1 ml LB in a microfuge tube and vortexed to resuspend cells. The resuspension was immediately plated on LB+50 μg/ml kanamycin. Resulting colonies represent transconjugants with the two antibiotic-selectable markers inserted at the *att*Tn7 site on the chromosome.

## Acknowledgements

We thank Ian Campbell, Karthik Hullahalli, and other members of the Waldor laboratory for helpful discussions and critical feedback on the manuscript. This article is subject to HHMI’s Open Access to Publications policy. HHMI laboratory heads have previously granted a non-exclusive CC BY 4.0 license to the public and a sublicensable license to HHMI in their research articles. Pursuant to those licenses, the author-accepted manuscript of this article can be made freely available under a CC BY 4.0 license immediately upon publication. This manuscript is the result of funding in whole or in part by the National Institutes of Health (NIH). It is subject to the NIH Public Access Policy. Through acceptance of this federal funding, NIH has been given a right to make this manuscript publicly available in PubMed Central upon the official date of publication, as defined by NIH.

## Data Availability

The *E. coli* MG1655 genomic reference sequence NC_000913.3 was used for mapping Tn-seq sample reads. Plasmid sequences will be made available on Addgene. Sequencing reads will be deposited in the Sequencing Read Archive (SRA).

## References

1. H. A. Shuman, T. J. Silhavy, The art and design of genetic screens: Escherichia coli. Nat Rev Genet 4, 419–431 (2003).

2. T. Van Opijnen, H. L. Levin, Transposon Insertion Sequencing, a Global Measure of Gene Function. Annu. Rev. Genet. 54, 337–365 (2020).

3. J. E. Peters, K. S. Makarova, S. Shmakov, E. V. Koonin, Recruitment of CRISPR-Cas systems by Tn7-like transposons. Proc. Natl. Acad. Sci. U.S.A. 114 (2017).

4. S. E. Klompe, P. L. H. Vo, T. S. Halpin-Healy, S. H. Sternberg, Transposon-encoded CRISPR–Cas systems direct RNA-guided DNA integration. Nature 571, 219–225 (2019).

5. J. Strecker, et al., RNA-guided DNA insertion with CRISPR-associated transposases. Science 365, 48– 53 (2019).

6. Y. Zhang, et al., Multicopy Chromosomal Integration Using CRISPR-Associated Transposases. ACS Synth. Biol. 9, 1998–2008 (2020).

7. P. L. H. Vo, et al., CRISPR RNA-guided integrases for high-efficiency, multiplexed bacterial genome engineering. Nat Biotechnol 39, 480–489 (2021).

8. S. Yang, et al., Orthogonal CRISPR-associated transposases for parallel and multiplexed chromosomal integration. Nucleic Acids Research 49, 10192–10202 (2021).

9. Z.-H. Cheng, et al., Repurposing CRISPR RNA-guided integrases system for one-step, efficient genomic integration of ultra-long DNA sequences. Nucleic Acids Research 50, 7739–7750 (2022).

10. B. E. Rubin, et al., Species- and site-specific genome editing in complex bacterial communities. Nat Microbiol 7, 34–47 (2021).

11. C. J. Tou, B. Orr, B. P. Kleinstiver, Precise cut-and-paste DNA insertion using engineered type V-K CRISPR-associated transposases. Nat Biotechnol 41, 968–979 (2023).

12. G. D. Lampe, et al., Targeted DNA integration in human cells without double-strand breaks using CRISPR-associated transposases. Nat Biotechnol 42, 87–98 (2024).

13. I. P. Witte, et al., Programmable gene insertion in human cells with a laboratory-evolved CRISPR-associated transposase. Science 388, eadt5199 (2025).

14. E. Aliu, K. Lee, K. Wang, CRISPR RNA-guided integrase enables high-efficiency targeted genome engineering in Agrobacterium tumefaciens. Plant Biotechnol J 20, 1916–1927 (2022).

15. P. Y. Pechenov, D. A. Garagulya, D. S. Stanovov, A. V. Letarov, New Effective Method of Lactococcus Genome Editing Using Guide RNA-Directed Transposition. IJMS 23, 13978 (2022).

16. L. Trujillo Rodríguez, A. J. Ellington, C. R. Reisch, M. G. Chevrette, CRISPR-Associated Transposase for Targeted Mutagenesis in Diverse Proteobacteria. ACS Synth. Biol. 12, 1989–2003 (2023).

17. Z. L. Yap, A. S. M. Z. Rahman, A. M. Hogan, D. B. Levin, S. T. Cardona, A CRISPR-Cas-associated transposon system for genome editing in Burkholderia cepacia complex species. Appl Environ Microbiol 90, e00699–24 (2024).

18. A. M. Garza Elizondo, J. Chappell, Targeted Transcriptional Activation Using a CRISPR-Associated Transposon System. ACS Synth. Biol. 13, 328–336 (2024).

19. K. M. Wetmore, et al., Rapid Quantification of Mutant Fitness in Diverse Bacteria by Sequencing Randomly Bar-Coded Transposons. mBio 6, 10.1128/mbio.00306-15 (2015).

20. W. Chen, et al., Targeted genetic screening in bacteria with a Cas12k-guided transposase. Cell Reports 36, 109635 (2021).

21. A. B. Banta, et al., A Targeted Genome-scale Overexpression Platform for Proteobacteria. bioRxiv 2024.03.01.582922 (2024). 10.1101/2024.03.01.582922.

22. M. C. Chao, S. Abel, B. M. Davis, M. K. Waldor, The design and analysis of transposon insertion sequencing experiments. Nat Rev Microbiol 14, 119–128 (2016).

23. D. Manna, S. Porwollik, M. McClelland, R. Tan, N. P. Higgins, Microarray analysis of Mu transposition in Salmonella enterica, serovar Typhimurium: transposon exclusion by high-density DNA binding proteins. Molecular Microbiology 66, 315–328 (2007).

24. S. Kimura, T. P. Hubbard, B. M. Davis, M. K. Waldor, The Nucleoid Binding Protein H-NS Biases Genome-Wide Transposon Insertion Landscapes. mBio 7, e01351–16 (2016).

25. D. Choe, et al., Revealing Causes for False-Positive and False-Negative Calling of Gene Essentiality in Escherichia coli Using Transposon Insertion Sequencing. mSystems 8, e00896–22 (2023).

26. G. D. Lampe, A. R. Liang, D. J. Zhang, I. S. Fernández, S. H. Sternberg, Structure-guided engineering of type I-F CASTs for targeted gene insertion in human cells. Nat Commun 16, 7891 (2025).

27. K. S. Makarova, E. V. Koonin, Annotation and Classification of CRISPR-Cas Systems. Methods Mol Biol 1311, 47–75 (2015).

28. T. S. Halpin-Healy, S. E. Klompe, S. H. Sternberg, I. S. Fernández, Structural basis of DNA targeting by a transposon-encoded CRISPR–Cas system. Nature 577, 271–274 (2020).

29. D. G. Gibson, et al., Enzymatic assembly of DNA molecules up to several hundred kilobases. Nat Methods 6, 343–345 (2009).

30. D. M. Stalker, R. Kolter, D. R. Helinski, Nucleotide sequence of the region of an origin of replication of the antibiotic resistance plasmid R6K. Proc. Natl. Acad. Sci. U.S.A. 76, 1150–1154 (1979).

31. L. Ferrières, et al., Silent Mischief: Bacteriophage Mu Insertions Contaminate Products of Escherichia coli Random Mutagenesis Performed Using Suicidal Transposon Delivery Plasmids Mobilized by Broad-Host-Range RP4 Conjugative Machinery. J Bacteriol 192, 6418–6427 (2010).

32. M. T. Petassi, S.-C. Hsieh, J. E. Peters, Guide RNA Categorization Enables Target Site Choice in Tn7-CRISPR-Cas Transposons. Cell 183, 1757-1771.e18 (2020).

33. E. R. Westra, et al., H-NS-mediated repression of CRISPR-based immunity in Escherichia coli K12 can be relieved by the transcription activator LeuO. Molecular Microbiology 77, 1380–1393 (2010).

34. D. C. Grainger, Structure and function of bacterial H-NS protein. Biochem Soc Trans 44, 1561–1569 (2016).

35. D. W. Basta, et al., Inducible transposon mutagenesis identifies bacterial fitness determinants during infection in mice. Nat Microbiol 10, 1171–1183 (2025).

36. B. Lang, et al., High-affinity DNA binding sites for H-NS provide a molecular basis for selective silencing within proteobacterial genomes. Nucleic Acids Research 35, 6330–6337 (2007).

37. D. F. Allison, G. G. Wang, R-loops: formation, function, and relevance to cell stress. Cell Stress 3, 38– 47 (2019).

38. T. Chen, C. Guestrin, XGBoost: A Scalable Tree Boosting System in Proceedings of the 22nd ACM SIGKDD International Conference on Knowledge Discovery and Data Mining, KDD ’16., (Association for Computing Machinery, 2016), pp. 785–794.

39. S. M. Lundberg, et al., From local explanations to global understanding with explainable AI for trees. Nat Mach Intell 2, 56–67 (2020).

40. K.-H. Choi, H. P. Schweizer, mini-Tn7 insertion in bacteria with single attTn7 sites: example Pseudomonas aeruginosa. Nat Protoc 1, 153–161 (2006).

41. A. B. Banta, R. A. Cuellar, N. Nadig, B. C. Davis, J. M. Peters, The promise of CRISPR-associated transposons for bacterial functional genomics. Current Opinion in Microbiology 83, 102563 (2025).

42. F. Tenjo-Castaño, S. S. Rout, S. Dey, G. Montoya, Unlocking the potential of CRISPR-associated transposons: from structural to functional insights. Trends in Genetics 41, 660–677 (2025).

43. M. W. G. Walker, S. E. Klompe, D. J. Zhang, S. H. Sternberg, Novel molecular requirements for CRISPR RNA-guided transposition. Nucleic Acids Research 51, 4519–4535 (2023).

44. L. C. T. Song, et al., Identification of Proteins Influencing CRISPR-Associated Transposases for Enhanced Genome Editing. [Preprint] (2024). Available at: https://www.biorxiv.org/content/10.1101/2024.09.11.612086v2 [Accessed 28 September 2025].

45. A. Roberts, M. A. Nethery, R. Barrangou, Functional characterization of diverse type I-F CRISPR-associated transposons. Nucleic Acids Research 50, 11670–11681 (2022).

46. S. G. Park, et al., Comprehensive profiling of activity and specificity of RNA-guided transposons reveals opportunities to engineer improved variants. Nucleic Acids Res 53, gkaf917 (2025).

47. B. P. Kleinstiver, et al., High-fidelity CRISPR–Cas9 nucleases with no detectable genome-wide off-target effects. Nature 529, 490–495 (2016).

48. M. W. Davis, E. M. Jorgensen, ApE, A Plasmid Editor: A Freely Available DNA Manipulation and Visualization Program. Front. Bioinform. 2 (2022).

49. S. E. Klompe, et al., Evolutionary and mechanistic diversity of Type I-F CRISPR-associated transposons. Molecular Cell 82, 616-628.e5 (2022).

50. C. L. Holmes, et al., Patterns of Klebsiella pneumoniae bacteremic dissemination from the lung. Nat Commun 16, 785 (2025).

51. J. M. Peters, et al., Enabling genetic analysis of diverse bacteria with Mobile-CRISPRi. Nat Microbiol 4, 244–250 (2019).

52. F. Pedregosa, et al., Scikit-learn: Machine Learning in Python. J. Mach. Learn. Res. 12, 2825–2830 (2011).

53. M. Tong, et al., Gene Dispensability in Escherichia coli Grown in Thirty Different Carbon Environments. mBio 11, e02259–20 (2020).

