## Supplementary Information for "Adapting CRISPR-associated transposons for rapid and high-throughput reverse genetics"

a

```
araD  1    ATGTTAGAAGATCTCAAACGCCAGGTATTAGAAGCCAACCTGGCGCTGCCAAAACACAAC  60
sgbE  1    ATGTTAGAGCAACTGAAAGCCGACGTGCTGGCGGCGAATCTGGCGCTTCCCGCTCACCAT  60

araD  61   CTGGTCACGCTCACATGGGGCAACGTCAGCGCCGTTGATCGCGAGCGCGGCGTCT--TTG  118
sgbE  61   CTGGTGACGTTCACCTGGGGCAATGTCAGCGCGGTAGA-CGAAA-CGCGGCAATGGATGG  118

araD  119  TGATCAAACCTTCCGGCGTCGATTACAGCGTCATGACCGCTGACGATATGGTCGTGGTTA  178
sgbE  119  TAATCAAACCTTCCGGCGTCGAGTACGACGTGATGACCGCCGACGATATGGTGGTGGTTG  178
                                                       6     12     18
araD  179  GCATCGAAACCGGTGAAGTGGTTGAAGGTACGAAAAAGCCCTCCTCCGACACGCCAACTC  238
sgbE  179  AGATAGCCAGCGGTAAGGTGGTGGAAGGCAGCAAAAAACCCTCTTCCGATACACCAACGC  238
           30
araD  239  ACCGGCTGCTCTATCAGGCATTCCCCTCCATTGGCGGCATTGTGCATACGCACTCGCGCC  298
sgbE  239  ATCTGGCGCTCTACCGTCGCTATGCCGAAATTGGCGGTATTGTGCATACCCACTCGCGCC  298

araD  299  ACGCCACCATCTGGGCGCAGGCGGGTCAGTCGATTCCAGCAACCGGCACCACCCACGCCG  358
sgbE  299  ACGCCACCATCTGGTCACAGGCCGGGCTGGATCTCCCCGCCTGGGGCACCACCCACGCCG  358

araD  359  ACTATTTCTACGGCACCATTCCCTGCACCCGCAAAATGACCGACGCAGAAATCAACGGCG  418
sgbE  359  ATTATTTTTACGGTGCCATCCCCTGCACGCGACAGATGACCGCAGAGGAGATTAACGGCG  418

araD  419  AATATGAGTGGGAAACCGGTAACGTCATCGTAGAAACCTTTGAAAAACAGGGTATCGATG  478
sgbE  419  AATATGAATATCAGACCGGCGAAGTGATCATTGAAACCTTCGAAGAACGTGGCAGGAGTC  478

araD  479  CAGCGCAAATGCCCGGC-GTTCTGGTCCATTCCCACGGCCCGTTTGCATGGGGCAAAAAT  537
sgbE  479  CGGCACAAAT-CCCGGCGGTGCTGGTGCATTCTCACGGCCCGTTCGCATGGGGTAAAAAC  537

araD  538  GCCGAAGATGCGGTGCATAACGCCATCGTGCTGGAAGAGGTCGCTTATATGGGGATATTC  597
sgbE  538  GCCGCCGATGCCGTGCATAACGCCGTAGTACTCGAAGAATGCGCCTATATGGGTCTATTC  597

araD  598  TGCCGTCAGTTAGCGCCGCAGTTACCGGATATGCAGCAAACG--CTGCTGGATAAACACT  655
sgbE  598  TCGCGCCAGCTTGCGCCGCAGCTCCCTGCGATGCA--AAACGAACTGCTGGATAAGCACT  655

araD  656  ATCTGCGTAAGCATGGCGCGAAGGCATATTACGGGCAGTAA  696
sgbE  656  ACCTGCGTAAGCATGGGGCCAATGCCTATTACGGGCAGTAA  696
```

Identities: 525/701 (75%), Gaps: 10/701 (1%)

b

```
                    12    18    24    30
araA CGGAAACCCTGCGTCAGGTCACCCAACATGCCGA
dgoT CCGTAACCCTGCGCCAGGTACCCCACCACCAGCG
```

c

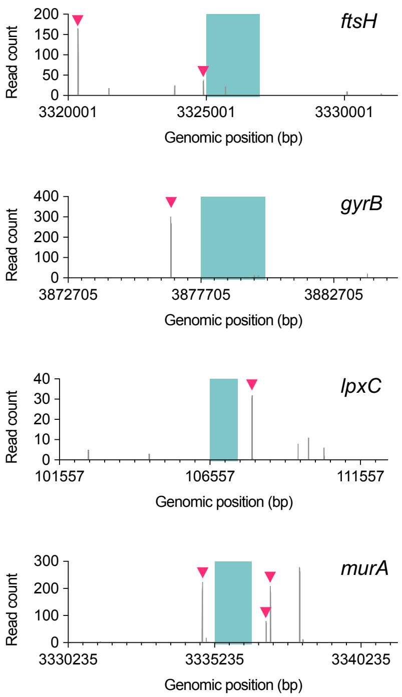

**Fig. S1. CAST is susceptible to off-target insertion. a)** Sequence alignment between the targeted gene *araD* and the untargeted gene *sgbE*. One guide designed for *araD* shares 25 of 32 nucleotides with a region in *sgbE*, despite lacking a canonical 'CN' PAM. The sixth position of the guide is dispensable for target recognition by Type I CRISPR systems. **b)** Sequence comparison between a guide targeting *araA* and its predicted off-target site in *dgoT*. **c)** Off-target insertions can occur when the distance between the guide's target site and the actual insertion site is unusually long. Such "missed" insertions may inflate estimates of mutant abundance, particularly for essential genes. The coding region of the essential gene is shaded in green. Red arrowheads indicate Read2-mapped insertions associated with Read1 guides targeting the nearby essential gene.

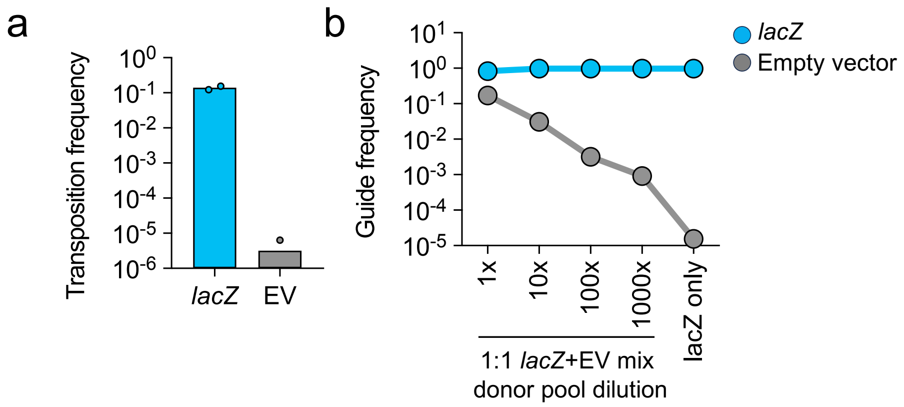

**Fig. S2. Independent confirmation of donor plasmid crosstalk. a)** Transposition frequency of a non-targeting donor plasmid (empty vector, EV) is >4 orders of magnitude lower than that of a *lacZ*-targeting donor, indicating minimal background activity. **b)** Relative guide frequency of *lacZ*-targeting and EV donor plasmids mixed 1:1 then serially diluted prior to conjugation. Dilution reduces the apparent EV guide frequency measured by PCR, indicating that donor crosstalk overestimates true insertion frequency.

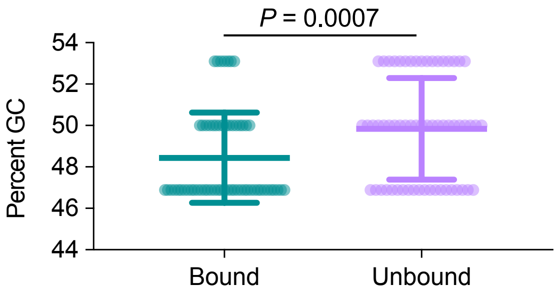

**Fig. S3. GC content of guides targeting H-NS bound and unbound regions.** Although the difference is statistically different, the average GC content is comparable between guides targeting H-NS-bound regions (48.5%) and those targeting unbound regions (49.8%). Mean ± S.D. is shown. *P* value was calculated using a one-sided Mann-Whitney test.

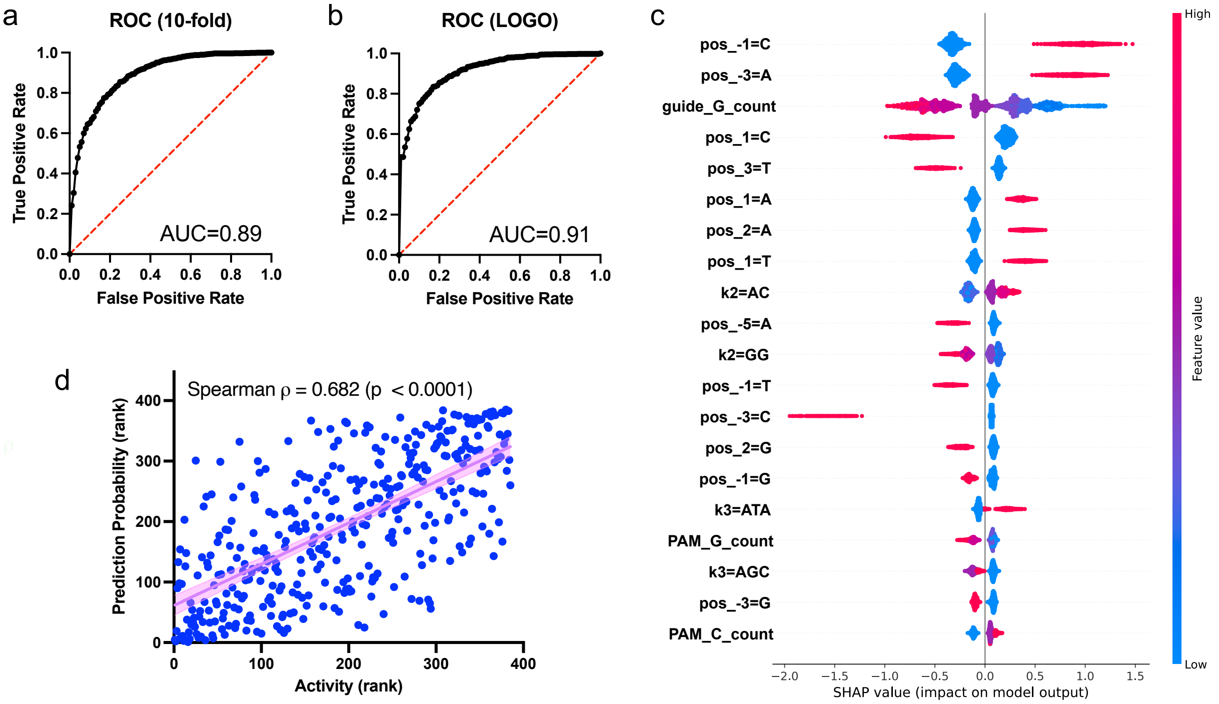

**Fig. S4. Internal validation of the predictive model and quantification of feature contributions.** Model performance under **a)** 10-fold cross validation and **b)** leave-one-gene-out (LOGO) validation. **c)** Beeswarm plot showing mean SHAP values for the top 20 sequence features most strongly associated with guide activity. **d)** Spearman correlation between observed guide activity and its prediction probability.

**Table S1. MultiCAST Pool1 internal validation (10-fold)**

| fold | n_train | n_test | AUROC | AUPRC | Precision | Recall | F1 |
|---|---|---|---|---|---|---|---|
| 1 | 2685 | 299 | 0.8630872 | 0.8601514 | 0.7439024 | 0.8133333 | 0.7770701 |
| 2 | 2685 | 299 | 0.8758837 | 0.8719454 | 0.8148148 | 0.7333333 | 0.7719298 |
| 3 | 2685 | 299 | 0.8887248 | 0.8748968 | 0.8297872 | 0.7852349 | 0.8068966 |
| 4 | 2685 | 299 | 0.8651007 | 0.8309714 | 0.7816901 | 0.7449664 | 0.7628866 |
| 5 | 2686 | 298 | 0.9055448 | 0.8970441 | 0.8367347 | 0.8255034 | 0.8310811 |
| 6 | 2686 | 298 | 0.9050944 | 0.8860173 | 0.8175676 | 0.8120805 | 0.8148148 |
| 7 | 2686 | 298 | 0.8845097 | 0.8836913 | 0.7616279 | 0.8791946 | 0.8161994 |
| 8 | 2686 | 298 | 0.896356 | 0.8926726 | 0.8150685 | 0.7986577 | 0.8067797 |
| 9 | 2686 | 298 | 0.8970767 | 0.8916902 | 0.8108108 | 0.8053691 | 0.8080808 |
| 10 | 2686 | 298 | 0.900545 | 0.893311 | 0.775641 | 0.8120805 | 0.7934426 |

**Table S2. MultiCAST Pool1 internal validation (leave-one-gene-out (LOGO))**

| heldout_gene | n_train | n_test | AUROC | AUPRC | Precision | Recall | F1 |
|---|---|---|---|---|---|---|---|
| acrA | 2939 | 45 | 0.943609 | 0.826832 | 0.3181818 | 1 | 0.482759 |
| acrB | 2930 | 54 | 0.945724 | 0.841906 | 0.6153846 | 1 | 0.761905 |
| ampG | 2909 | 75 | 0.849673 | 0.656254 | 0.5714286 | 0.83333 | 0.677966 |
| araA | 2920 | 64 | 0.951136 | 0.977636 | 0.9722222 | 0.79545 | 0.875 |
| araB | 2905 | 79 | 0.912899 | 0.934305 | 0.9411765 | 0.68085 | 0.790123 |
| araC | 2944 | 40 | 0.905371 | 0.868893 | 0.7142857 | 0.88235 | 0.789474 |
| araD | 2963 | 21 | 0.888889 | 0.983312 | 1 | 0.55556 | 0.714286 |
| araE | 2929 | 55 | 0.936937 | 0.845568 | 0.6666667 | 1 | 0.8 |
| argA | 2914 | 70 | 0.892992 | 0.93863 | 0.9714286 | 0.70833 | 0.819277 |
| argB | 2955 | 29 | 0.847826 | 0.955394 | 0.8888889 | 0.69565 | 0.780488 |
| argC | 2945 | 39 | 1 | 1 | 1 | 0.6875 | 0.814815 |
| argE | 2936 | 48 | 0.938889 | 0.958297 | 0.9565217 | 0.73333 | 0.830189 |
| argG | 2918 | 66 | 0.929101 | 0.970834 | 0.9069767 | 0.86667 | 0.886364 |
| argH | 2923 | 61 | 0.962567 | 0.985389 | 0.9487179 | 0.84091 | 0.891566 |
| cysC | 2958 | 26 | 0.83125 | 0.879299 | 0.8235294 | 0.875 | 0.848485 |
| cysD | 2955 | 29 | 0.789216 | 0.786605 | 0.6153846 | 0.66667 | 0.64 |
| cysG | 2928 | 56 | 0.865625 | 0.770734 | 0.6315789 | 0.75 | 0.685714 |
| cysH | 2956 | 28 | 0.909091 | 0.727498 | 0.5454545 | 1 | 0.705882 |
| cysI | 2920 | 64 | 0.907843 | 0.916166 | 0.8181818 | 0.9 | 0.857143 |
| cysJ | 2913 | 71 | 0.909468 | 0.84855 | 0.71875 | 0.82143 | 0.766667 |
| cysN | 2919 | 65 | 0.923295 | 0.920524 | 0.8709677 | 0.81818 | 0.84375 |
| fruA | 2924 | 60 | 0.919312 | 0.811181 | 0.625 | 0.83333 | 0.714286 |
| fruB | 2946 | 38 | 0.955182 | 0.956479 | 0.9411765 | 0.7619 | 0.842105 |
| fruK | 2944 | 40 | 0.919271 | 0.916825 | 0.625 | 0.9375 | 0.75 |
| galK | 2936 | 48 | 0.927734 | 0.86362 | 0.5555556 | 0.9375 | 0.697674 |
| galM | 2934 | 50 | 0.977778 | 0.751429 | 0.2777778 | 1 | 0.434783 |
| galP | 2937 | 47 | 0.89819 | 0.69218 | 0.5714286 | 0.92308 | 0.705882 |
| galR | 2949 | 35 | 0.894558 | 0.831984 | 0.6875 | 0.78571 | 0.733333 |
| galT | 2937 | 47 | 0.87013 | 0.755159 | 0.6111111 | 0.78571 | 0.6875 |
| hisA | 2965 | 19 | 0.888889 | 0.91308 | 0.8888889 | 0.8 | 0.842105 |
| hisB | 2939 | 45 | 0.942387 | 0.930813 | 0.7272727 | 0.88889 | 0.8 |
| hisC | 2939 | 45 | 0.96 | 0.964388 | 0.9047619 | 0.76 | 0.826087 |
| hisD | 2930 | 54 | 0.907433 | 0.878837 | 0.7894737 | 0.65217 | 0.714286 |
| hisF | 2955 | 29 | 1 | 1 | 0.625 | 1 | 0.769231 |
| hisG | 2953 | 31 | 0.850877 | 0.744847 | 0.7272727 | 0.66667 | 0.695652 |
| hisI | 2965 | 19 | 0.869048 | 0.640523 | 0.6 | 0.85714 | 0.705882 |
| ilvA | 2936 | 48 | 0.96 | 0.960555 | 0.95 | 0.76 | 0.844444 |

| | | | | | | | |
|---|---|---|---|---|---|---|---|
| ilvC | 2916 | 68 | 0.967777 | 0.987793 | 1 | 0.63265 | 0.775 |
| ilvD | 2913 | 71 | 0.901961 | 0.956137 | 0.893617 | 0.82353 | 0.857143 |
| ilvE | 2949 | 35 | 0.893939 | 0.948406 | 0.8571429 | 0.75 | 0.8 |
| lacI | 2950 | 34 | 0.973333 | 0.945674 | 0.6428571 | 1 | 0.782609 |
| lacZ | 2907 | 77 | 0.868673 | 0.826907 | 0.6666667 | 0.88235 | 0.759494 |
| leuA | 2909 | 75 | 0.920058 | 0.949398 | 0.9705882 | 0.76744 | 0.857143 |
| leuB | 2947 | 37 | 0.773529 | 0.805406 | 0.8333333 | 0.5 | 0.625 |
| leuC | 2932 | 52 | 0.93702 | 0.960322 | 0.8928571 | 0.80645 | 0.847458 |
| leuD | 2961 | 23 | 0.969231 | 0.981335 | 1 | 0.92308 | 0.96 |
| metA | 2951 | 33 | 0.916667 | 0.932479 | 0.9047619 | 0.90476 | 0.904762 |
| metB | 2930 | 54 | 0.953846 | 0.982423 | 0.969697 | 0.82051 | 0.888889 |
| metC | 2929 | 55 | 0.946779 | 0.967803 | 0.8857143 | 0.91176 | 0.898551 |
| metE | 2902 | 82 | 0.897581 | 0.960921 | 0.9555556 | 0.69355 | 0.803738 |
| metF | 2943 | 41 | 0.941799 | 0.97043 | 0.9545455 | 0.77778 | 0.857143 |
| metL | 2908 | 76 | 0.906154 | 0.940586 | 0.8974359 | 0.7 | 0.786517 |
| metR | 2939 | 45 | 0.875 | 0.947691 | 0.9545455 | 0.65625 | 0.777778 |
| pflA | 2962 | 22 | 0.717949 | 0.685744 | 0.5 | 0.77778 | 0.608696 |
| proA | 2935 | 49 | 0.875 | 0.794065 | 0.6363636 | 0.82353 | 0.717949 |
| proB | 2947 | 37 | 0.896825 | 0.838519 | 0.5833333 | 0.77778 | 0.666667 |
| proC | 2959 | 25 | 1 | 1 | 0.4285714 | 1 | 0.6 |
| serA | 2929 | 55 | 0.978723 | 0.878292 | 0.3333333 | 1 | 0.5 |
| serB | 2952 | 32 | 0.94686 | 0.909453 | 0.5714286 | 0.88889 | 0.695652 |
| thrA | 2902 | 82 | 0.922101 | 0.942758 | 0.8536585 | 0.76087 | 0.804598 |
| thrB | 2941 | 43 | 0.960591 | 0.983084 | 1 | 0.86207 | 0.925926 |
| thrC | 2943 | 41 | 0.937799 | 0.951771 | 0.8888889 | 0.72727 | 0.8 |

**Table S3. MultiCAST Pool3 external validation**

| AUROC | AUPRC | Precision | Recall | F1 |
| --- | --- | --- | --- | --- |
| 0.871526525 | 0.99221203 | 0.9835391 | 0.7331288 | 0.8400703 |

**Table S4. Primer sequences for Tn-seq and pooled guide amplification**

**Tn-seq PCR primers**

| Name | Sequence |
|---|---|
| CASTnseq-fwd | AATGATACGGCGACCACCGAGATCTACACTCTTTCCCTACACGACGCTCTTCCGATCTTCATTACTACTGCAAAGTAGCTGATAAC |
| P7-AD01-index-Rev | CAAGCAGAAGACGGCATACGAGATCGTGATGTGACTGGAGTTCAGACGTGTG |
| P7-AD02-index-Rev | CAAGCAGAAGACGGCATACGAGATACATCGGTGACTGGAGTTCAGACGTGTG |
| P7-AD03-index-Rev | CAAGCAGAAGACGGCATACGAGATGCCTAAGTGACTGGAGTTCAGACGTGTG |
| P7-AD04-index-Rev | CAAGCAGAAGACGGCATACGAGATTGGTCAGTGACTGGAGTTCAGACGTGTG |

**Tn-seq oligo adapters**

| Name | Sequence |
|---|---|
| Fork truncated-NH2 | TACCACGACCA-NH2 |
| Index Fork-rev | GTGACTGGAGTTCAGACGTGTGCTCTTCCGATCTGGTCGTGGTAT |

**Pooled guides PCR primers**

| Name | Sequence |
|---|---|
| Var21 | AATGATACGGCGACCACCGAGATCTACACTCTTTCCCTACACGACGCTCTTCCGATCTAATGATGGGTTAAAAAGGATCGATCC |
| Var22 | AATGATACGGCGACCACCGAGATCTACACTCTTTCCCTACACGACGCTCTTCCGATCTATGCGATGGGTTAAAAAGGATCGATCC |
| Var23 | AATGATACGGCGACCACCGAGATCTACACTCTTTCCCTACACGACGCTCTTCCGATCTTGCACGATGGGTTAAAAAGGATCGATCC |
| Var24 | AATGATACGGCGACCACCGAGATCTACACTCTTTCCCTACACGACGCTCTTCCGATCTTCATTCGATGGGTTAAAAAGGATCGATCC |
| Var25 | AATGATACGGCGACCACCGAGATCTACACTCTTTCCCTACACGACGCTCTTCCGATCTGAATCGAGATGGGTTAAAAAGGATCGATCC |
| Var26 | AATGATACGGCGACCACCGAGATCTACACTCTTTCCCTACACGACGCTCTTCCGATCTGTCAACTTGATGGGTTAAAAAGGATCGATCC |
| Var27 | AATGATACGGCGACCACCGAGATCTACACTCTTTCCCTACACGACGCTCTTCCGATCTCGGCGTGGCGATGGGTTAAAAAGGATCGATCC |
| Var28 | AATGATACGGCGACCACCGAGATCTACACTCTTTCCCTACACGACGCTCTTCCGATCTCCTGTACCTTGATGGGTTAAAAAGGATCGATCC |
| AD001 | CAAGCAGAAGACGGCATACGAGATCGTGATGTGACTGGAGTTCAGACGTGTGCTCTTCCGATC |
| AD002 | CAAGCAGAAGACGGCATACGAGATACATCGGTGACTGGAGTTCAGACGTGTGCTCTTCCGATC |
| AD003 | CAAGCAGAAGACGGCATACGAGATGCCTAAGTGACTGGAGTTCAGACGTGTGCTCTTCCGATC |
| AD004 | CAAGCAGAAGACGGCATACGAGATTGGTCAGTGACTGGAGTTCAGACGTGTGCTCTTCCGATC |

| | |
|---|---|
| AD005 | CAAGCAGAAGACGGCATACGAGATCACTGTGTGACTGGAGTTCAGACGTGTGCTCTTCCGATC |
| AD006 | CAAGCAGAAGACGGCATACGAGATATTGGCGTGACTGGAGTTCAGACGTGTGCTCTTCCGATC |
| AD007 | CAAGCAGAAGACGGCATACGAGATGATCTGGTGACTGGAGTTCAGACGTGTGCTCTTCCGATC |
| AD008 | CAAGCAGAAGACGGCATACGAGATTCAAGTGTGACTGGAGTTCAGACGTGTGCTCTTCCGATC |
| AD009 | CAAGCAGAAGACGGCATACGAGATCTGATCGTGACTGGAGTTCAGACGTGTGCTCTTCCGATC |
| AD010 | CAAGCAGAAGACGGCATACGAGATAAGCTAGTGACTGGAGTTCAGACGTGTGCTCTTCCGATC |
| AD011 | CAAGCAGAAGACGGCATACGAGATGTAGCCGTGACTGGAGTTCAGACGTGTGCTCTTCCGATC |
| AD012 | CAAGCAGAAGACGGCATACGAGATTACAAGGTGACTGGAGTTCAGACGTGTGCTCTTCCGATC |
| AD013 | CAAGCAGAAGACGGCATACGAGATTTGACTGTGACTGGAGTTCAGACGTGTGCTCTTCCGATC |
| AD014 | CAAGCAGAAGACGGCATACGAGATGGAACTGTGACTGGAGTTCAGACGTGTGCTCTTCCGATC |
| AD015 | CAAGCAGAAGACGGCATACGAGATTGACATGTGACTGGAGTTCAGACGTGTGCTCTTCCGATC |
| AD016 | CAAGCAGAAGACGGCATACGAGATGGACGGGTGACTGGAGTTCAGACGTGTGCTCTTCCGATC |
| AD018 | CAAGCAGAAGACGGCATACGAGATGCGGACGTGACTGGAGTTCAGACGTGTGCTCTTCCGATC |
| AD019 | CAAGCAGAAGACGGCATACGAGATTTTCACGTGACTGGAGTTCAGACGTGTGCTCTTCCGATC |
| AD020 | CAAGCAGAAGACGGCATACGAGATGGCCACGTGACTGGAGTTCAGACGTGTGCTCTTCCGATC |
| AD021 | CAAGCAGAAGACGGCATACGAGATCGAAACGTGACTGGAGTTCAGACGTGTGCTCTTCCGATC |
| AD022 | CAAGCAGAAGACGGCATACGAGATCGTACGGTGACTGGAGTTCAGACGTGTGCTCTTCCGATC |
| AD023 | CAAGCAGAAGACGGCATACGAGATCCACTCGTGACTGGAGTTCAGACGTGTGCTCTTCCGATC |
| AD025 | CAAGCAGAAGACGGCATACGAGATATCAGTGTGACTGGAGTTCAGACGTGTGCTCTTCCGATC |
| AD027 | CAAGCAGAAGACGGCATACGAGATAGGAATGTGACTGGAGTTCAGACGTGTGCTCTTCCGATC |
